# SARS-CoV-2 NSP1 C-terminal region (residues 130-180) is an intrinsically disordered region

**DOI:** 10.1101/2020.09.10.290932

**Authors:** Amit Kumar, Ankur Kumar, Prateek Kumar, Neha Garg, Rajanish Giri

## Abstract

Nonstructural protein 1 (NSP1) of SARS-CoV-2 plays a key role in downregulation of RIG-I pathways and interacts with 40 S ribosome. Recently, the cryo-EM structure in complex with 40S ribosome is deciphered. However, the structure of full length NSP1 without any partner has not been studies. Also, the conformation of NSP1-C terminal region in isolation is not been studied. In this study, we have investigated the conformational dynamics of NSP1C-terminal region (NSP1-CTR; amino acids 130-180) in isolation and under different solvent environments. The NSP1-CTR is found to be intrinsically disordered in aqueous solution. Further, we used alpha helix inducer, trifluoroethanol, and found induction of alpha helical conformation using CD spectroscopy. Additionally, in the presence of SDS, NSP1-CTR is showing a conformational change from disordered to ordered, possibly gaining alpha helix in part. But in presence of neutral lipid DOPC, a slight change in conformation is observed. This implies the possible role of hydrophobic interaction and electrostatic interaction on the conformational changes of NSP1. The changes in structural conformation were further studied by fluorescence-based studies, which showed significant blue shift and fluorescence quenching in the presence of SDS and TFE. Lipid vesicles also showed fluorescence-based quenching. In agreement to these result, fluorescence lifetime and fluorescence anisotropy decay suggests a change in conformational dynamics. The zeta potential studies further validated that the conformational dynamics is mostly because of hydrophobic interaction. In last, these experimental studies were complemented through Molecular Dynamics (MD) simulation which have also shown a good correlation and testify our experiments. We believe that the intrinsically disordered nature of the NSP1-CTR will have implications in disorder based binding promiscuity with its interacting proteins.

## 1. Introduction

Among four genera of coronaviruses (CoVs), alpha and beta are able to infect humans. CoVs have single-stranded positive-sense RNA genome, of which, the two-third part (~20kb) is translated into two large polyproteins named pp1a and pp1ab. These two polyproteins encode sixteen non-structural proteins (NSP1–NSP16), which make up the viral replication-transcription complex (1, 2). The very interesting feature among the four CoVs genera is presence of NSP1, which is present only in alpha- and beta-genera of CoVs (3). This feature may correlate with the pathogenicity of CoV as alpha and beta are highly pathogenic than the gamma and delta genera (3). The NMR structure of SARS-CoV Nsp1 showed that N- and the C-terminal domains are disordered with β-barrel fold in between (4). However, it is stated that the SARS-CoV NSP1 does not possess any enzymatic activity. The NSP1 have been found to downregulate the immune function of host cells through RIG-1 signaling pathway (5, 6). Also, NSP1 bound with the 40S ribosomal subunit and degrade the host mRNA (6, 7). SARS-CoV-2 NSP1 along with NSP3, NSP12, NSP13, NSP14, ORF3, ORF6 and M protein are found to inhibit the Sendai virus-induced IFN-β promoter activation (8). Moreover, the NSP1 size varies among different lineage of beta-CoVs, and share different degree of amino acid sequence similarity (3, 6). Despite of the different amino acid composition, the high structural similarity of NSP1 found to regulate the same feature in all genera of alpha and beta-CoV (9).

For a period of time bioinformatics studies show the penetrance of IDPs in viruses and all domain of life from bacteria to eukaryotes (10–14). Recently, bioinformatics based study by our group showed that NSP1 of SARS-CoV-2 regions (amino acids 137-145 and 172-179) are identified as a molecular recognition feature (MoRFs) region (2). Generally, most of IDPs are unfolded in solution and switch to a conformation when bind with particular ligands and undergo coupled folding and binding (15–18). The IDPs capabilities to participate in multiple cellular signaling lies in their structural amenability, which can be relate to the presence of MoRFs and their kinetics advantages (19–23). Therefore, the molecular interactions of IDPs are fugitive and dynamic, which allow them to compete and exchange the binding partners in function regulating proteins. Given the fact that the NSP1 majorly affect the host protein, and intracellular environment is highly crowded (19, 24, 25). Therefore, the impact of surrounding environment on the conformational behavior of NSP1-CTR need to be addressed to assess its disorder-function paradigm, which eventually lead us to understand the interaction mechanistic with respective partners. Furthermore, given the earlier prediction that the C-terminal region of NSP1 has molecular recognition features, so it is important to investigate the folding propensity in the presence of lipids. We should ask these questions in terms of an IDP folding behavior. It is important as what conformational changes this NSP1 CTR can adopt upon interaction with synthetic membrane environments such as liposomes? What about the conformational propensity in presence of organic solvents? What is the impact of temperature on this peptide? To answer these questions, in the present study, structural conformations of SARS-CoV-2 NSP1 with respect to C-terminal disordered regions is explored in different experimental condition, which gives a deeper insight to structural conformation in respective environment.

## 2. Material and methods

### 2.1 Chemicals and reagents

The NSP1 peptide (residues 130–180), corresponding to the C-terminal regions with purity>82% was purchased from Genescript, USA. Organic solvents such as Trifluoroethanol (TFE) with ≥99% purity was purchased from Sigma-Aldrich. Lyophilized peptide was dissolved in ultra-pure water at a concentration of 1 mg/ml.

### 2.2 Phylogenetic analysis of SARS-CoV-2 on the basis of NSP1

The full length and aligned sequences fasta files were obtained after blastp search. The phylogenetic tree was constructed to locate the taxonomic position of the SARS-CoV-2 NSP1 using MEGA analysis tool v5. Further, the NSP1 (50 residues) were studied with Clustal Omega tool to reveal their conserved sequences and represented with the help of ESPript 3.0.

### 2.3 Secondary structural analysis of NSP1 and insight into intrinsic disorder properties

The sequence of NSP1 C-terminal (residues 130-180) “NH2-AGGHSYGADLKSFDLGDELGTDPYEDFQENWNTKHSSGVTRELMRELNGG-COOH” was retrieved from the UniProt (ID: P0DTC1.1). As described earlier by our group, protein intrinsic disorder predictor, PONDR^®^ VSL2 was used for analysis of intrinsic disorder properties of NSP1-CTR (2, 13, 14, 19, 26, 27). The secondary structure predisposition analysis for NSP1-CTR was performed with several different web-servers pep2d, Jpred4, and PSIPRED (28–30).

### 2.4 Liposome preparation

The liposome were prepared as described earlier (19). Briefly, the neutral lipid DOPC (dioleoyl-phosphatidyl-ethanolamine) and negatively charged lipid DOPS (1,2-dioleoyl-sn-glycero-3-phospho-Lserine) were purchased from Avanti Polar Lipids (Alabaster, AL). The chloroform from the lipid solution was removed using rotary evaporator at 40°C and the dry lipid films were hydrated in 50 mM phosphate buffer (pH 7.4). The final concentration of the DOPC and DOPS liposomes were 37.09 mM and 24.69 mM, respectively. The resulting suspension was freeze-thaw-vortex in liquid nitrogen and water at 60 °C, following which the lipids were extruded 25 times through the mini extruder (Avanti Polar Lipids, Inc. USA) through cut off filter of 100nm polycarbonate membrane to prepare uniform Large unilamellar vesicles (LUV). The size of LUVs was determined by dynamic light scattering using Zetasizer Nano ZS (Malvern Instruments Ltd., UK). The size of DOPS and DOPC was 125 nm and 102 nm, respectively.

### 2.5 Circular Dichroism spectroscopy

Circular dichroism spectrometer MOS500 (BioLogic) and JASCO machine (Jasco J-1500 CD spectrometer) was used for CD data recording. 20 μM peptide sample were prepared in 50 mM phosphate buffer, pH 7.0. Peptide was kept in organic solvents (TFE) with increasing concentration from 0 to 90% and far-UV (190–240 nm) spectra were recorded in 1 mm quartz cuvette. Similarly, the peptide was assessed for structural changes in Sodium Dodecyl Sulfate (SDS) (below and above critical micellar concentration (CMC)) and in the presence of DOPS and DOPC LUVs (pH 7.4) All the spectra were recorded at scan speed of 50 nm/min with a response time of 1s and 1 nm bandwidth. The equivalent spectra of buffers were recorded and subtracted from the spectra of the test samples. The ellipticity with HT value of less than 600 volt were considered to plot the spectra.

### 2.6 Phase diagram analysis

Another method called “phase analysis” was utilized to study the intermediates qualitatively from conformational transitions under different conditions as described earlier (19, 20). This method involves the analysis of spectral data by building the phase diagram for spectral intensities I (λ1) versus I (λ2) where I is the spectral intensity measured at wavelengths λ1 and λ2 under different experimental conditions.

### 2.7 Fluorescence spectroscopy

We monitored the intrinsic Trp fluorescence intensity in NSP1. A 5μM peptide in 50mM sodium phosphate buffer (pH 7.0) was prepared with increasing concentration of TFE and SDS, and 20 μM peptide (pH 7.4) in DOPS and DOPC. Emission spectra was recorded from 300 to 500 nm at 295nm excitation wavelength in a Horiba Fluorolog-3 spectrofluorometer. The individual negative blank was subtracted from each test samples (20).

### 2.8 Fluorescence lifetime measurement

Fluorescence lifetime measurements were carried out for the 5μM peptide (pH 7.0) in increasing concentration of TFE and SDS, and 20 μM peptide (pH 7.4) in DOPC and DOPS by using Fluorolog TCSPC (Horiba Scientific Inc.) with pulsed LED sources. The excitation wavelength was set at 284 nm, and the emission wavelength was kept at 345nm. A Ludox solution was used for correct instrument response factor (IRF). Fluorescence decay data was analyzed through Data Analysis Software provided by Horiba Scientific. Decay data was fitted tri-exponentially decay function with a best fit chi-squared value approaching 1.0 in order to calculate the fluorescence lifetimes.

### 2.9 Time-Resolved Fluorescence Anisotropy Decay

Time-Resolved Fluorescence Anisotropy Decay were carried out for the 30 μM peptide (pH 7.4) in increasing concentration of SDS by using Fluorolog TCSPC (Horiba Scientific Inc.) with pulsed LED sources. The measurement range of 200 ns, peak preset of 1000 counts was set up to monitor the decay with emission wavelength of 345 nm. Ludox was used to monitor the prompt decay (at 284 nm wavelength) to correct the instrument response factor (IRF). Fluorescence anisotropy measures the depolarization, which happens because the energy transfer to another molecule of different orientation. Alternatively, molecular rotation is also caused by Brownian motion and the local environment i.e., viscosity, size of molecule, and molecular confinement affect the molecular motion (31). Thus, a measurement of fluorescence anisotropy is useful in obtaining information concerning molecular size and mobility.

The time-resolved fluorescence anisotropy decay function, r(t) is given as:

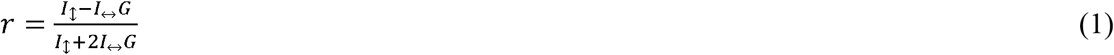

Where, *I*_↕_ and *I*_↔_ designate fluorescence decays obtained for parallel and perpendicular emission polarizations, respectively. Where G is the instrumental correction factor at the wavelength λ of emission and is the ratio of efficiency of detection for vertically and horizontally polarized light.

The anisotropy decay curve can be expressed by the following equation

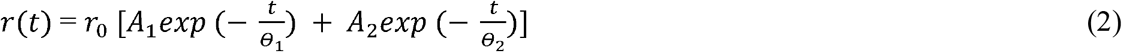

Where, *r*(*t*) is the anisotropy at time *t, r*_0_ is the initial anisotropy and ranges from 0.4 (parallel transition dipoles) to - 0.2 (perpendicular dipoles), *A*_1_ and *A*_2_ are the amplitude associated with the rotational correlation time *θ*_1_ and *θ*_2_, respectively.

### 2.10 Zeta potential measurement

NSP1 C-terminal interaction with DOPS and DOPC was studied for zeta potential measurements using Malvern Zetasizer Nano ZS device. Briefly, 200 μM of liposomes incubated with different concentration of NSP1 peptide at 25°C and measured for charge in zeta potential. A total of 3 measurements with 50 runs each were obtained using disposable zeta cells with platinum gold-coated electrodes (Malvern). The electrophoretic mobility obtained was used for the zeta potential calculation with the help of the Smoluchowski equation.

### 2.11 Molecular Dynamics (MD) Simulation of NSP1 C-terminal model

The atomic-level dynamic and energetics could be understood through computer simulation for appropriate time periods. Long simulation are helpful to explore structural properties and behavior with respect to time (32). We have utilized Pep-Fold (33, 34) webserver for constructing a 3D model for C-terminal of NSP1. It applies a coarse-grained (CG) forcefield and performs up to 200 simulation runs to build an energy minimized structure. The resultant model was then prepared using Chimera by addition of missing hydrogens and proper parameterization of asymmetrical residues (35).

We used Gromacs v5, where simulation setup was built by placing the protein structure in a cubic box along with SPC water model, 0.15M NaCl salt concentration. After solvation, the system was charge neutralized with counterions. To attain an energy minimized simulation system, the steepest descent method was used until the system was converged within 1000 kJ/mol. Further, the equilibration of system was done to optimize solvent in the environment. Using NVT and NPT ensembles within periodic boundary conditions for 100ps each, the system was equilibrated. The average temperature at 300K and pressure at 1 bar were maintained using Nose-Hoover and Parrinello-Rahman coupling methods during simulation. All bond-related constraints were solved using SHAKE algorithm. The final production run was performed for 500ns in our high performing cluster at IIT Mandi. After analyzing the trajectory, the last frame (we named ‘NSP1-F1’ to avoid confusion) was chosen for further studies in different conditions.

Next, the structural conformations of C-terminal of NSP1 (NSP1-F1) were analyzed in two different conditions of solvents: TFE (Trifluoroethanol; 8M) and SDS (Sodium dodecyl sulfate) mixed with water in separate simulation runs. By using above described forcefield parameters and addition of TFE and SDS molecules into the system, the simulation was performed for 300ns each. Further to investigate the structural transitions upon lipid-mediated environments, we performed molecular dynamics simulation of frame NSP1-F1 in presence of DOPS. Using previously described parameters for simulation in DOPS (36), we used CHARMM-GUI web interface (37) and inserted 300 molecules of DOPS around protein structure. Following minimization and equilibration, the final production MD was performed for 200ns. All trajectory analysis, calculations were performed using Chimera, maestro and Gromacs commands for calculating root mean square deviation (RMSD), root mean square fluctuation (RMSF), radius of gyration (Rg) for protein structure compactness, and solvent accessible surface area (SASA) for C-α atoms. The secondary structure component percentage was calculated using 2struc web server (38).

## 3. Results and discussion

### 3.1 Phylogenetic and Structural analysis of NSP1 C-terminal and insight into intrinsic disorder properties

Using neighbor-joining method, the phylogenetic tree of coronaviruses was constructed for fulllength amino acid sequences of NSP1 (180 residues) **(Figure 1A)**. It was clear from the multiple sequence alignment that Bat-SARS-CoV, SARS CoV and SARS-CoV-2 share consensus sequences. The SARS-CoV-2 C-terminal has few diverse amino acid residues, which indicated their evolutionary divergence from other coronaviruses **(Figure 1B)**.

**Figure 1.**
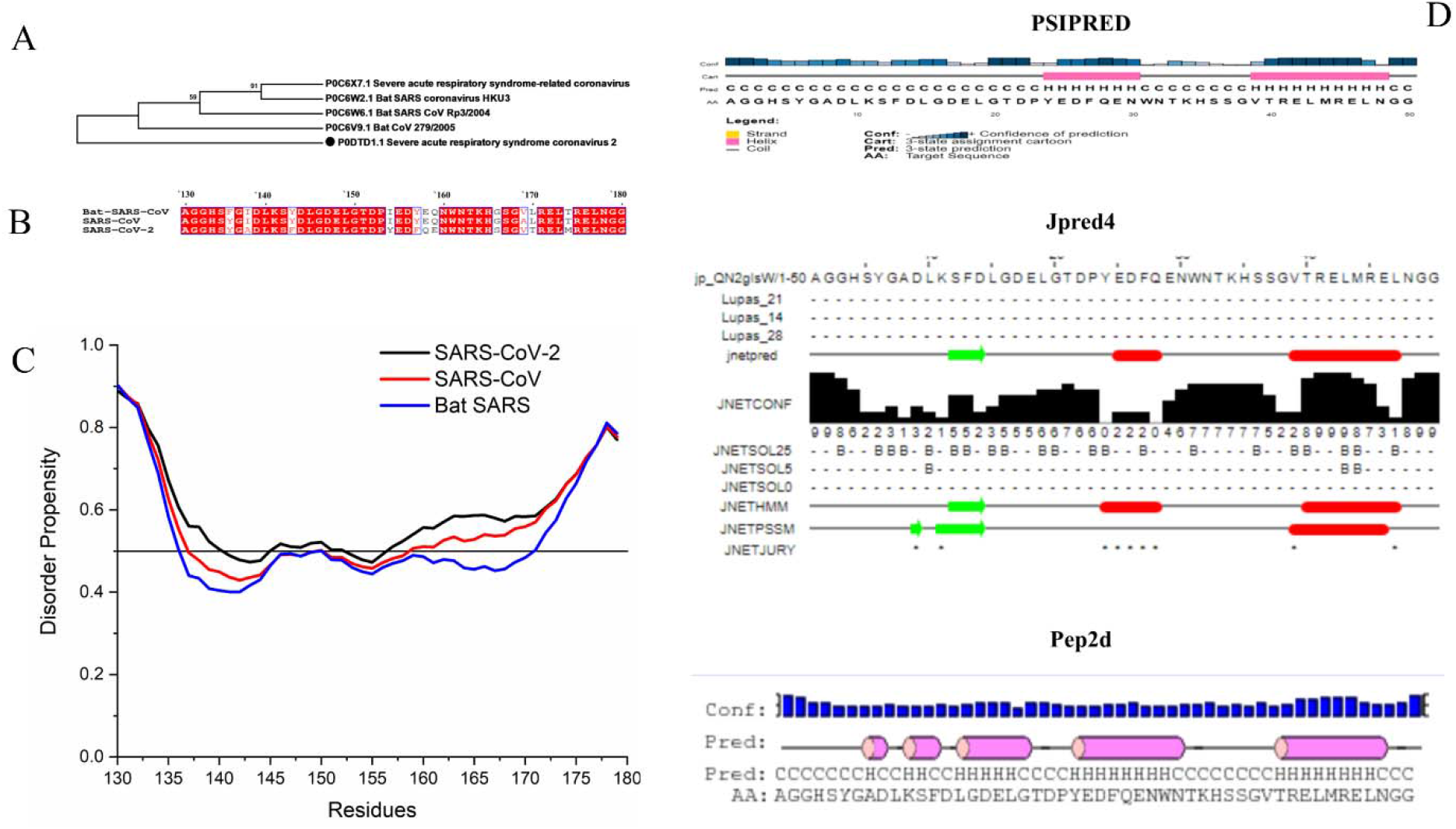
**(A)** The evolutionary history was inferred using the Neighbor-Joining method by using NSP1 full-length sequence. The percentage of replicate trees in which the associated taxa clustered together in the bootstrap test (500 replicates) are shown next to the branches. All positions containing gaps and missing data were eliminated. Evolutionary analyses were conducted in MEGA5. **(B)** The multiple sequence alignment of NSP1-CTR showing consensus sequences. **(C)** Disorder predisposition predictions of the NSP1-CTR (50 residues) using disorder predictor PONDR^®^ VSL2. The interception at 0.5 score on Y-axis shows the cut-off of disorderedness. Regions above than 0.5 are considered as disordered. **(D)** Secondary structure predisposition predictions using servers PSIPRED, Jpred4, and Pep2D (C-Coil, E-Extended strand, B-Beta strand, and H-Helix).

The intrinsic disorder predisposition of NSP1 C-terminal was studied using commonly used VSL2 predictor of intrinsic disorder and found that it is on borderline between ordered and disordered nature, near the prediction cutoff of 0.5 for disorder prediction (**Figure 1C).** Further, sequence-based secondary structure analysis was done by utilizing multiple webservers, which clearly showed that the part of C-terminal can acquire helical conformation. The PSIPRED has predicted a potential for helical structure at the 154-160 and 169-178 region **(Figure 1D).** Whereas, Jpred4 has indicated the β-strand propensity at 141-144 region and helical propensity at155-158, and 169-177 regions. While, Pep2d predicted the very high α-helical propensity at multiple regions, which was 141-142, 145-149, 154-161, and 169-178 regions **(Figure 1D)**.

### 3.2. NSP1-CTR is intrinsically disordered in isolation and has folding propensity in presence of organic solvents and lipids

At a concentration of 20 μM in phosphate buffer (pH 7.0), the CD spectrum of NSP1-CTR was dominated by a single minimum (199 nm), which indicates that this region in isolation acquire random coil/disordered conformation **(Figure 2A)**. To investigate the folding propensity of the disordered proteins organic solvents and lipids are generally used (39). Therefore, in this study, we have used SDS, TFE and liposomes for their hydrophobic/hydrophilic interface and charge mimic properties. CD Spectra at high SDS concentration showed the negative ellipticities at 208 and 222 nm, which is a typical characteristic of alpha-helical conformation. The effect of SDS on conformation of NSP1-CTR was observed and found that SDS in its monomeric form (0.5 mM) doesn’t have significant effect on conformation of NSP1. However, SDS above critical micellar concentration (CMC; 5mM and 50 mM) induce the helicity **(Figure 2B)**. Further, TFE was also found to induce helicity in NSP1-CTR disorder system **(Figure 2D)**. The secondary structure analysis was performed using K2D3 indicates the gain in alpha helical structure and loss in random coil in the presence of SDS and TFE **(Figure 2 C, E, F).** While, in the presence of DOPS and DOPC there is partial change in the structural conformation which we can’t say that its alpha helical and neither disordered **(Figure 2 G, H, I)**. The SDS micelles provide a model for the hydrophobic and hydrophilic interface similar to lipid membranes (40). Studies showed that the water make hydrogen bond with carbonyl group for a time and then water oxygen start to pair with N-H, and thus disrupt the intramolecular hydrogen bonding (41). Whereas, TFE has low dielectric constant and high dipole moment compared to water. The TFE directly interact with peptide and imparts hydrophobic effect. Irrespective to water, TFE make stable hydrogen bonding with carbonyl group of peptides and does not disrupt the intramolecular hydrogen bonding (41). In conclusion, there is change in secondary structure conformation in the presence of SDS, TFE and DOPC. Our finding is in accordance with the previous disorder system, where SDS, TFE, and liposomes were used as model system to mimic the biological environment such as hydrophobic nature, and protein-lipids interaction (19, 20, 36).

**Figure 2.**
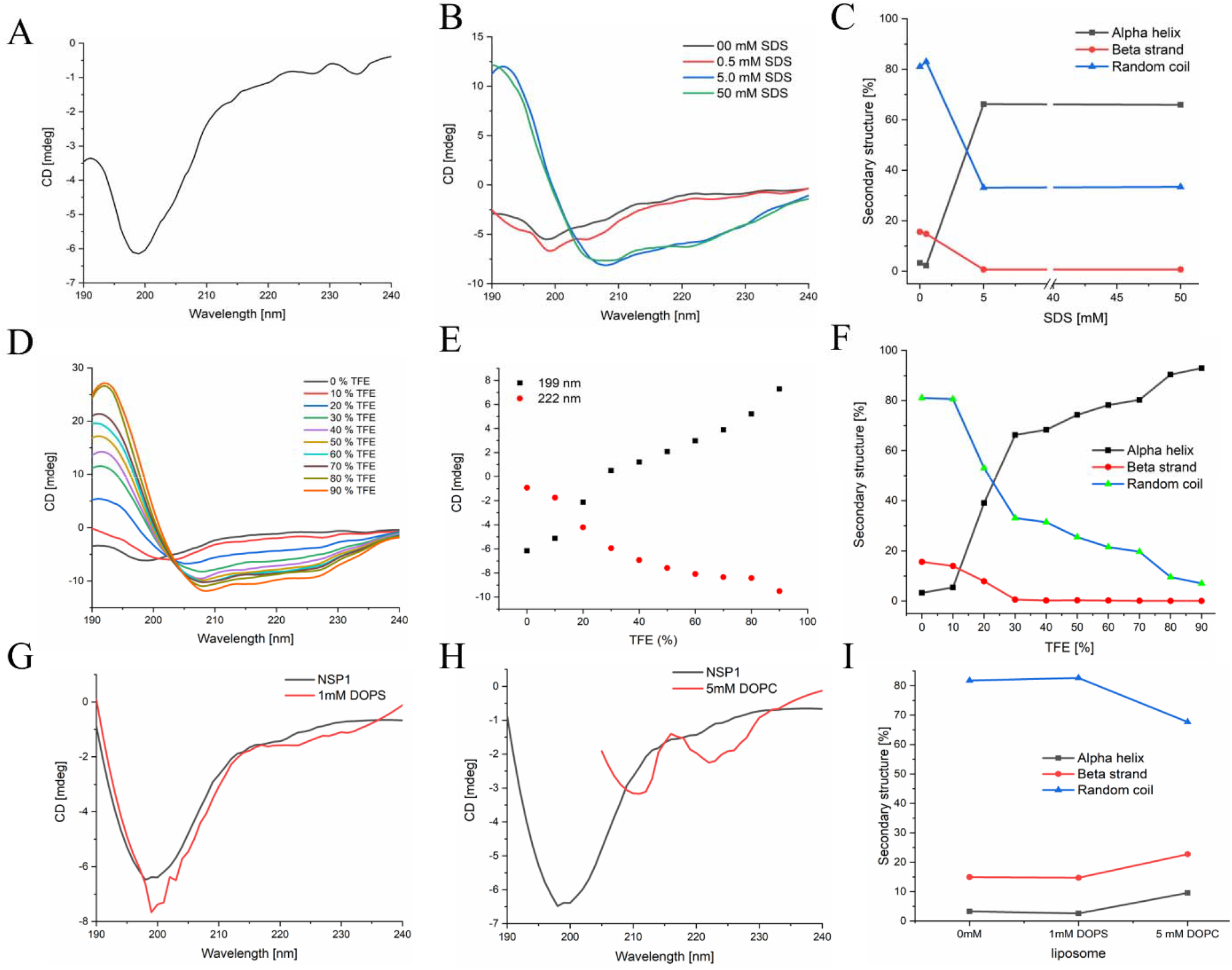
CD spectra showing effect of varying conditions on NSP1 C-terminal (A) 50 mM phosphate buffer (pH 7.0) (B) SDS micelle (D) TFE (G) DOPS and (H) DOPC. Panel (E) showing phase diagram change in structural conformational in the presence of TFE. Panel (C, F, I) showing the secondary structure analysis of NSP1 in SDS, TFE and liposomes, respectively.

Further, the validation of experimental results was done through computational simulation. Firstly, the model built through PEP-FOLD server was simulated in presence of SPC water model using Gromos54A7 forcefield for 500ns. Initially, the model constituted three helices (residues _145_LGDEL_149_, _155_EDFQE_159_, and _169_VTRELMREL_177_) mainly at C-terminal. After first simulation, the NSP1-CTR model started losing its structure and lost significant amount of helix; designated as NSP1-F1 **(Figure 3)**. The region with unstable helix was found to be 145-149 while other two helices had insignificant fluctuations. As calculated using an online server *2struc*, the simulated structure has lost its 10% helicity from 42% to 32%, as shown in **Figure 3.** This indicates that the NSP1-C-terminal has less structure and more disorder character in isolation.

**Figure 3:**
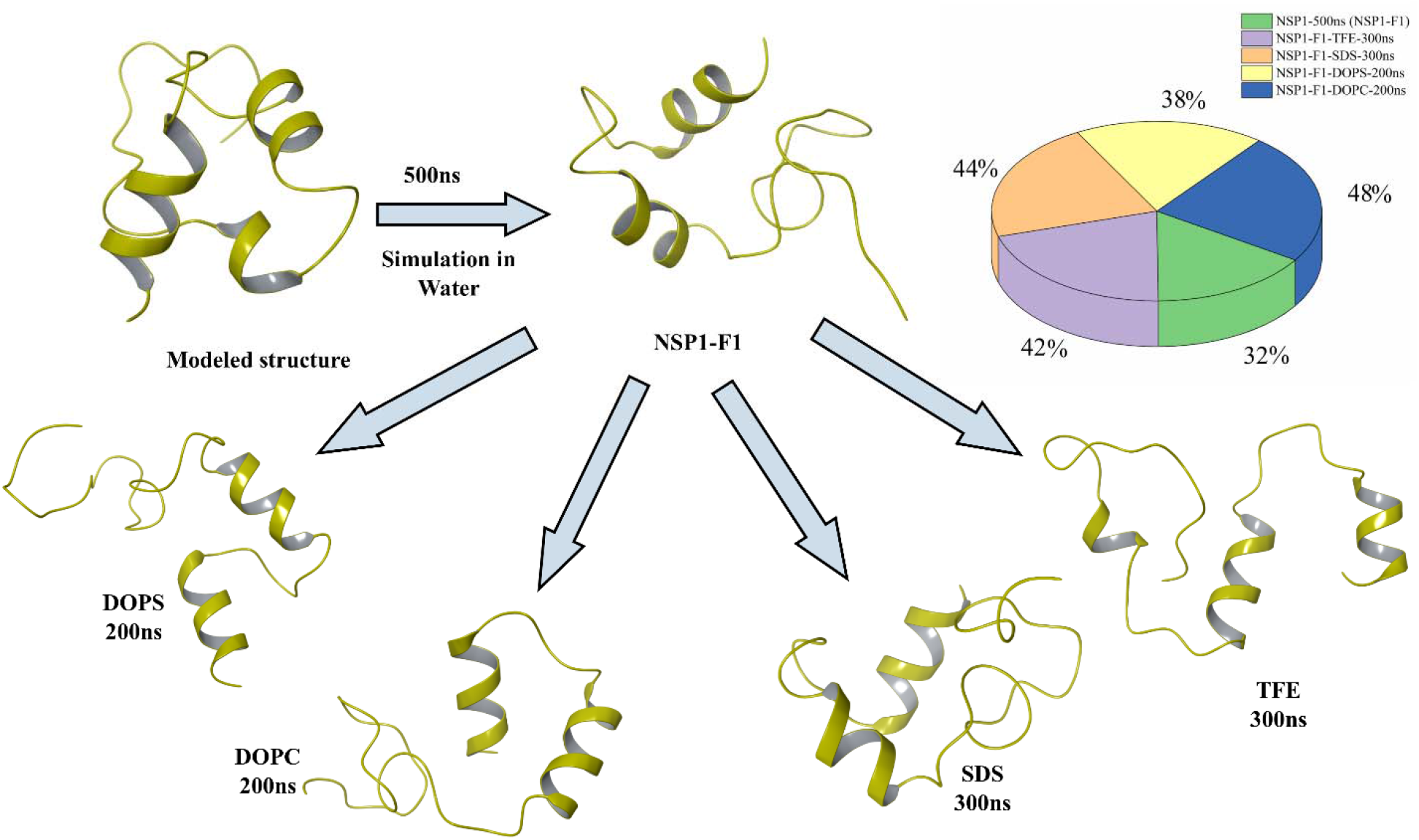
MD Simulation of modeled structure in water and its last frame trajectory in SDS, TFE, DOPC and DOPS. In inset, pie chart representing helix percentage in modeled structure and frames of NSP1 obtained from MD simulation.

The simulated model at 500 ns trajectory frame (NSP1-F1), was used for further simulation to study its ability to gain of structure. The first physiological condition was provided with 8M trifluoroethanol in the simulation box and the rest of the box was filled by water molecules. In presence of organic solvent, the NSP1-F1 structure showed increased helicity after 100ns up to 38% from 32% **(Figure 3)**. Subsequently, after 300ns, the trajectory frame shows 42% of helix present in the structure at residues 145-148, 155-163, and 171-177. Here also, the MD simulation results are purely correlated with the experimental observations and suggest that in organic solvent environment like TFE, the C-terminal of NSP1 has propensity to gain structure. Previously, simulation studies of amyloid beta-peptide showed that the peptide conformation changes at different solutions of TFE (27%, 40%, 60%, and 70%) (42). Moreover, a model peptide and an antimicrobial peptide were also demonstrated in co-presence of TFE with water, where the helical propensity of the structure was observed to be increased (43).

Further, MD simulation of NSP1-F1was performed in presence of SDS and DOPC and DOPS. In 200ns simulation time and in presence of nearly 60 molecules of SDS, the simulated frame or NSP1-F1 structure gained a slight increment in helical conformation at residues 155-161 and 170-178. Similarly, DOPS has also showed slight change in helical conformation at residues 155-162 and 170-177. In case of DOPC lipid environment, the residues 155-164 and 170-178 have shown helical conformation. As calculated by *2struc* server, the simulated structure in DOPS and DOPC has gained helicity from 32% to 38% and 48%, respectively, as shown in **Figure 3.** A similar work by Jalili et al., that has performed all atoms and coarse-grained simulation of Amyloid-beta (Aβ40) peptide in presence of SDS and have shown the formation of micelles around the peptide due to self-assembly of molecules of SDS (44).

### 3.3. Effect of temperature on the conformation of NSP1-CTR

Temperature attribute significant impact on the protein’s backbone. It is stated that the increase in the temperature leads to the change in secondary structure of intrinsically disordered proteins, which is associated with the hydrophobic interactions (19, 20). As depicted in **Figure 4A**, increase in temperature leads to partial change in structural conformation of NSP1-CTR. Further, to know the impact of temperature on NSP1-CTR and lipid association, effect of increasing temperature was studied **(Figure 4 B,C)**. DOPC induced more contraction to this IDP system as compared to the contraction without the lipids. The secondary structure analysis using K2D3 also suggested change in conformation in the presence of DOPC **(Figure 4D, E, F)**. The partial compaction was because of collapse in hydrophobic interaction with respect to secondary structure formation, however in the presence of DOPC changes in structural conformation was stable (19, 45). This suggest that NSP1-CTR interact with the neutral charged liposome and acquire stable conformation, which in isolation was disordered. Previously, non-helical region and hydrophilic chains in proteins was also found responsible for the contraction at high temperature (46, 47). DOPS somehow doesn’t enhance the contraction because both the peptide and lipid are negatively charged and they may not be binding together.

**Figure 4.**
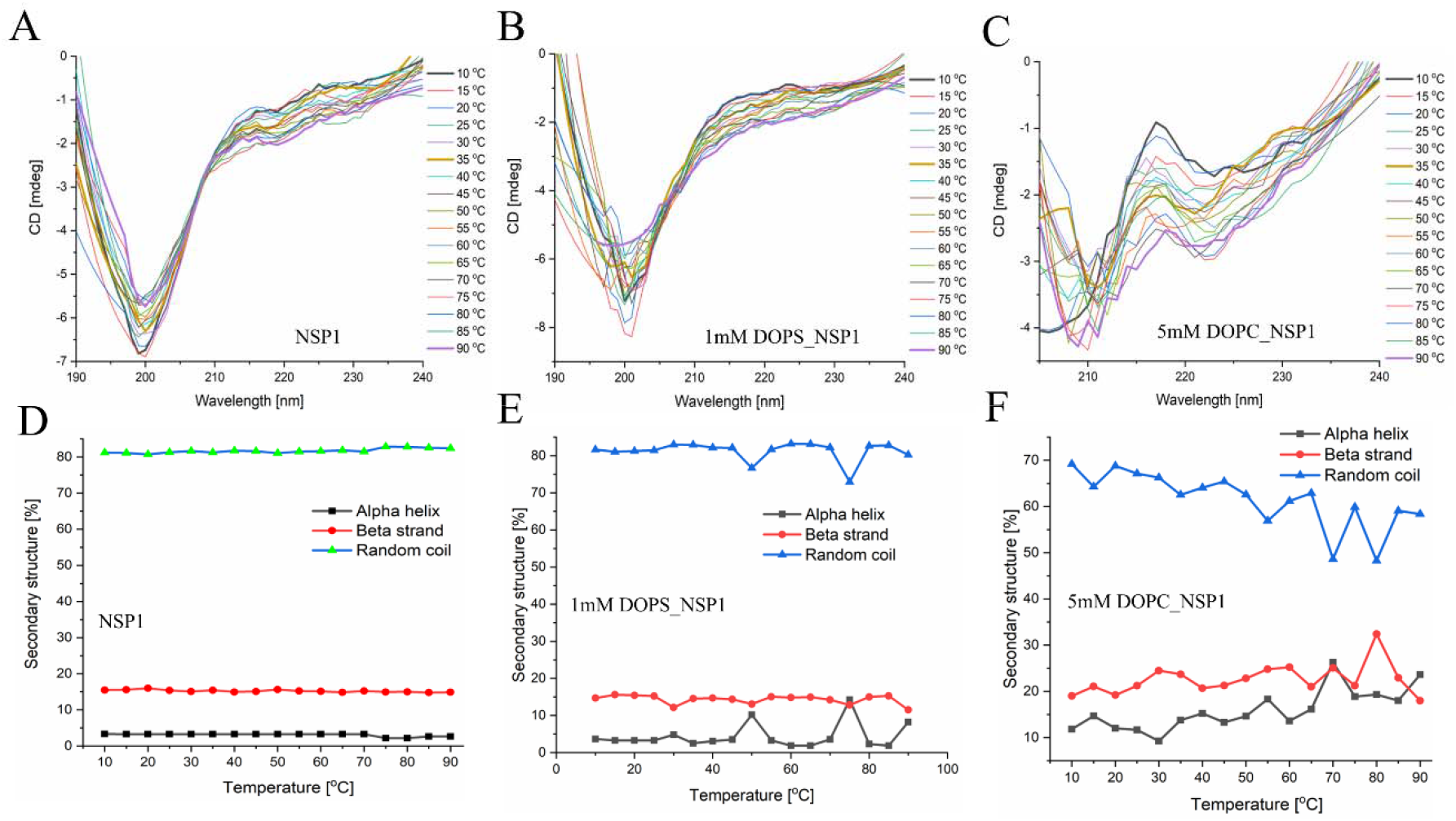
Effect of temperature on NSP1-CTR. Panel (A, D) showing the change in conformation and secondary structure in the presence of increasing temperature. Panel (B, E) and (C, F) showing the change in conformation and secondary structure in the presence of DOPS and DOPC, respectively.

### 3.4. NSP1-CTR conformational change revealed by Trp fluorescence

The NSP1-CTR has one Trp^161^ which showed emission maxima at 346 nm in phosphate buffer. With a rising concentration of SDS (above CMC), we observed decreased fluorescence intensity and a significant blue shift (emission maxima at 335nm) **(Figure 5A)**. The change in the fluorescence intensity of NSP1-CTR is due to conformational dynamics which leads Trp residue to expose to the SDS environment. Similarly, the presence of TFE also decreases the fluorescence intensity with blue shift (emission maxima at 339 nm) that may be due to increased non-polarity of the solvent **(Figure 5B)**. Further, NSP1-CTD interaction with the liposome (negatively charged liposome: DOPS; and neutral charged liposome: DOPC) also reduces the fluorescence intensity with blue shift (emission maxima at 345 nm and 344 nm for DOPS and DOPC, respectively; see **Figure 5C**). So, fluorescence studies suggest that the SDS, TFE, and liposome induces a conformational change in NSP1-CTR by imparting hydrophobic interaction and altering the hydrogen bonding. This fluorescence-based study was further validated by the lifetime measurement of the Trp fluorescence. A fluorophore (Trp^161^ in our case) in response to excitation spends time in the excited state, emit a photon and return to the ground state, which is denoted as its fluorescence lifetime. Any change in the physicochemical conditions leads to the change in the property of fluorophore and thus the fluorescence lifetime. The average lifetime of fluorophore Trp in NSP1-CTR was observed to be 1.84 ns in phosphate buffer (pH 7.0), while 1.02 ns in the presence of 5 mM SDS, and 1.57 ns in 60 % TFE **(Table 1** and **Figure 5D, E)**. The average lifetime in the presence of DOPS and DOPC decreased to 0.634 ns and 0.0847 ns, respectively **(Table 1** and **Figure 5F)**. Therefore, it can be seen that the lifetime decreases gradually in the presence of TFE. However, there is a sharp decrease of lifetime in the presence of SDS and liposome (both DOPC and DOPS).

**Figure 5.**
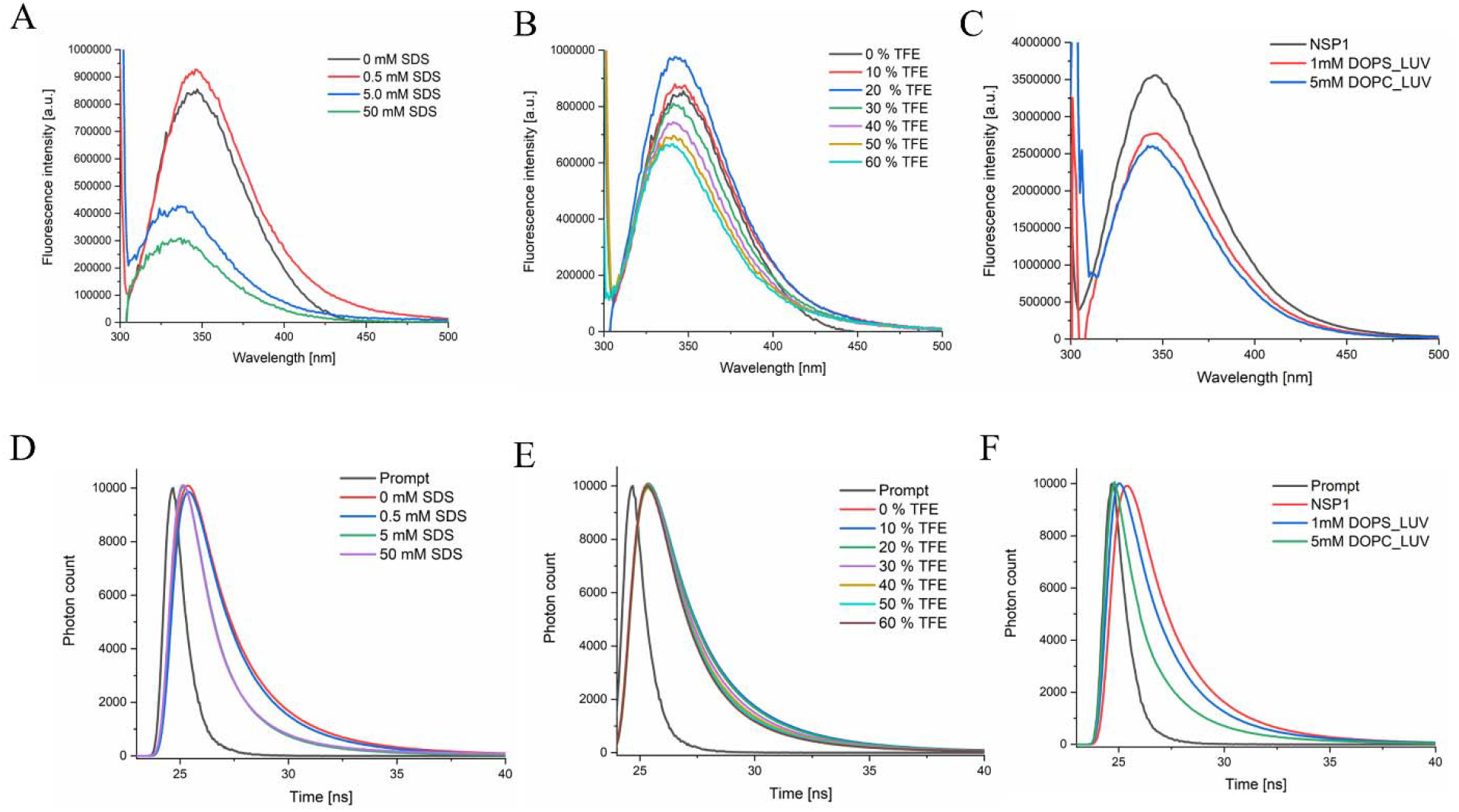
Tryptophan emission fluorescence analysis and a comparison of fluorescence lifetime decay curve of NSP1-CTR peptide in (A, D) SDS and (B, E) TFE and (C, F) Liposomes. In panel D the decay trace is of 0 mM and 0.5 mM of SDS is overlapping.

**Table 1.** Average lifetime in different conditions after fitting data in three exponential term. In case of SDS and TFE the 5 μM peptide concentration is considered. For Liposomes, 20 μM peptide concentration is considered.

| Component | Concentration | Average lifetime (Nanosecond) | CHISEQ ($\chi^2$) |
| --- | --- | --- | --- |
| NSP1-CTR only | 5 μM | 1.84 | 1.054 |
| SDS | 0.5 mM | 1.73 | 1.109 |
|  | 5 mM | 1.02 | 1.159 |
|  | 50 mM | 1.02 | 1.345 |
| TFE | 10 % | 1.91 | 1.154 |
|  | 20 % | 1.83 | 1.112 |
|  | 30 % | 1.72 | 1.097 |
|  | 40 % | 1.62 | 1.156 |
|  | 50 % | 1.61 | 1.152 |
|  | 60 % | 1.57 | 1.061 |
| Liposome | 1mM DOPS | .634 | 1.289 |
|  | 5 mM DOPC | .0847 | 1.088 |

Further, we have analyzed the NSP1-CTR conformational dynamics using Time-resolved fluorescence anisotropy decay measurements, which have been used previously for other IDPs (48, 49). The faster rotational correlation times (θ_1_) suggest a local motion of the fluorophore, whereas slower correlation times (θ_2_) is designated with segmental dynamics of peptide backbone and global tumbling of the entire peptide (31, 49). We measured the conformational dynamics and steric hindrance around the NSP1-CTD using Trp^161^ fluorophore in the SDS and organic solvent environment. **Figure 6** shows the fluorescence anisotropy decay for the NSP1-CTR in the presence of SDS micelle, organic solvents like TFE and ethanol. The decay curve was fitted with the bi-exponential decay function that can be correlated with the equation 2 as described in the method section. The rotational correlation time θ_1_ (faster rotational correlation time), was observed to be around 1 ns in all conditions **(Table 2).** Here the θ_1_ represents the the rotational correlation time of Trp^161^ residue motion. However, rotational correlation time θ_2_ (slower rotational correlation time) was observed to be beyond 200 ns at higher concentration of SDS, TFE and ethanol, represents the global conformational dynamics of NSP1-CTR. The slower rotational correlation times (θ_2_) for NSP1-CTR only is 16. 94 ns but it reduces to 4.09 ns, and 7.3 ns in 20% TFE and 20 % ethanol, respectively. Further, in the presence of SDS (5mM and 50 mM) and TFE (60%) where helical conformation was observed (monitored by CD; see **Figure 2)**, NSP1-CTR are showing θ_2_ between 1300-1600 ns. At higher concentration of SDS or TFE, a very long θ_2_ (between 1300-1600 ns) beyond 200 ns of measurement range represents very slow global rotation of NSP1-CTR. At 20 % TFE the lower value of θ_2_ as compared to higher concentration of TFE (60 %) and in the absence of TFE may suggest an existance of a transition phase where partially folded NSP1-CTD dominated. The slow rotation in case of SDS may be due to the self aggregation nature of SDS forms micellar assemblies that increases the size **(Table 2**). A similar type of approach has been employed previously to investigate the conformational dynamics of α-synuclein in the presence of SDS micelle using NMR that shows chemical entity of micelle doesn’t perturbed the conformation change but quantitative binding does (52). Amphiphilic nature of SDS molecules allows them to self-aggregate and form micellar assemblies (50–52). Overall, we may say that NSP1-CTR can acquire different structural conformation to perform some functional roles in host cells, which need further experimental validation. Previously some evidence suggests that the two helical structures, alpha 1 and alpha 2 (Trp^161^ lies in between these two helical regions) acquired by NSP1-CTR when interacted with the 40S ribosome subunit (6). Thus, our study corroborates these results and found a change in structural conformation. Recently, our group has demonstrated that p53 TAD2 was an intrinsically disordered protein and evidenced a similar finding with an organic solvent, which relates its diverse conformations with different interacting partners (20).

**Figure 6.**
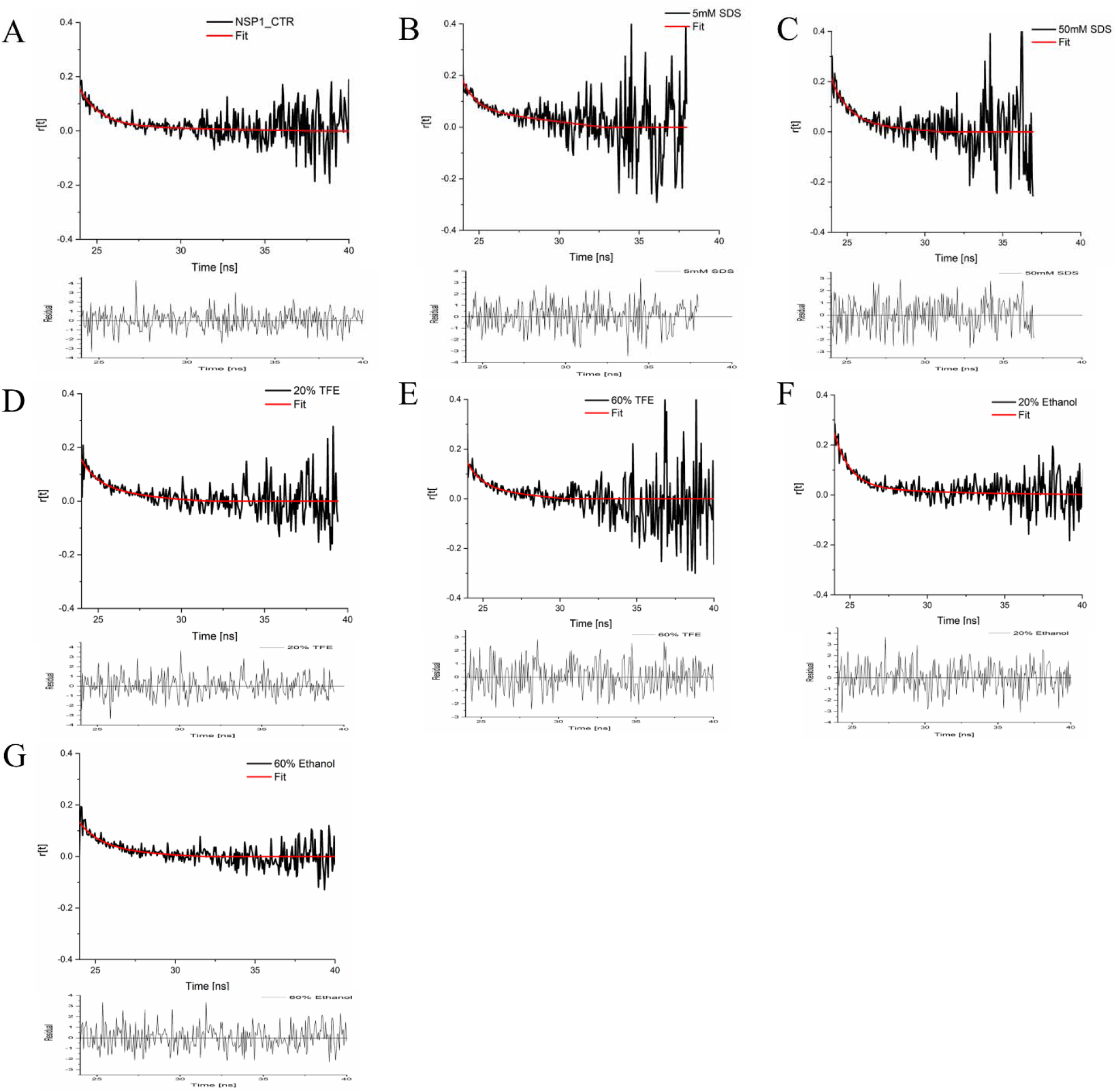
Decay of the fluorescence anisotropy of NSP1-CTR in the presence of (A) Buffer (B,C) SDS (D,E) TFE and (F,G) Ethanol.

**Table 2.**
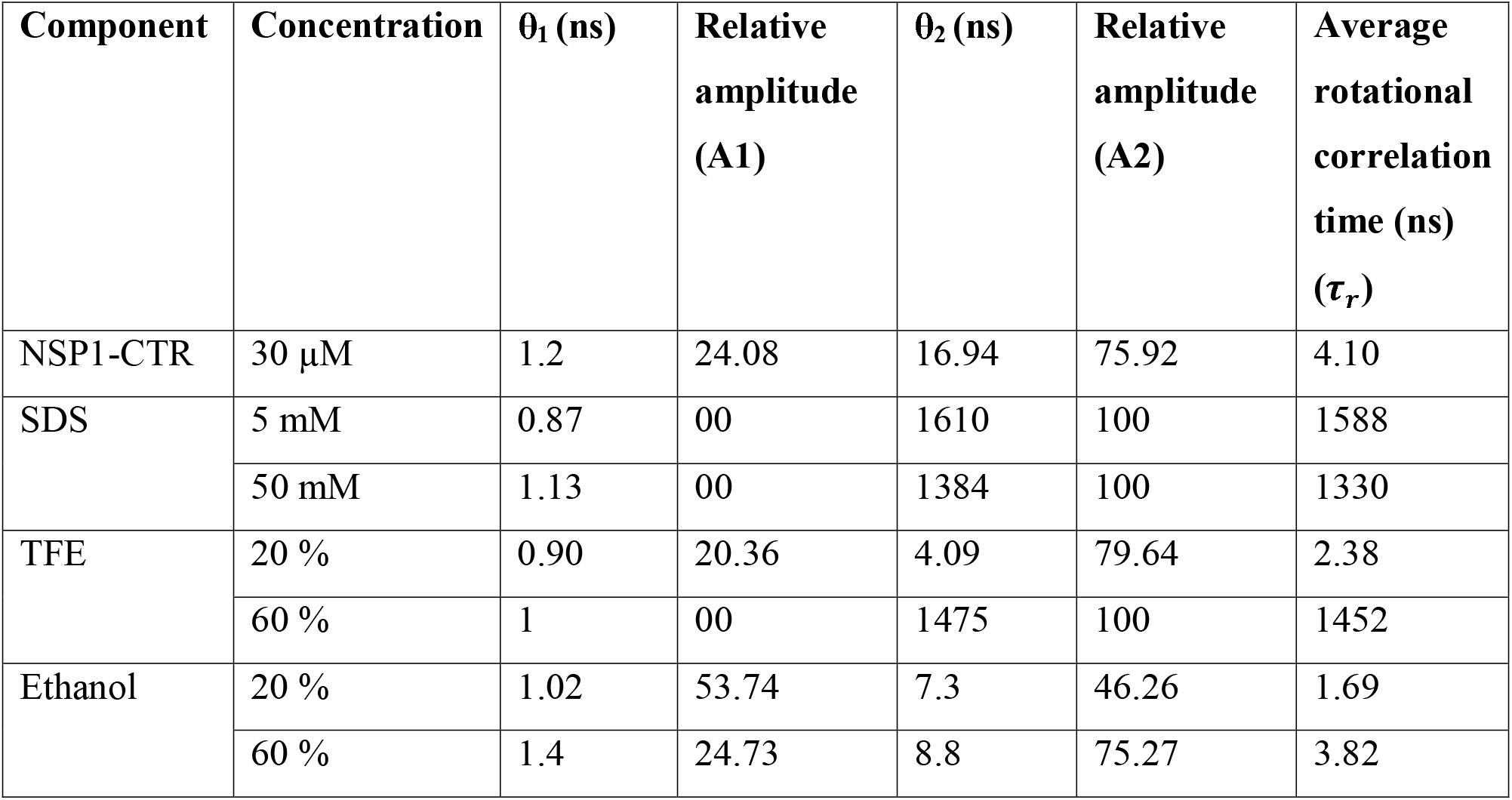
Anisotropy Decay Parameters for NSP1-CTR.

| Component | Concentration | $\theta_1$ (ns) | Relative amplitude (A1) | $\theta_2$ (ns) | Relative amplitude (A2) | Average rotational correlation time (ns) ($\tau_r$) |
| --- | --- | --- | --- | --- | --- | --- |
| NSP1-CTR | 30 μM | 1.2 | 24.08 | 16.94 | 75.92 | 4.10 |
| SDS | 5 mM | 0.87 | 00 | 1610 | 100 | 1588 |
|  | 50 mM | 1.13 | 00 | 1384 | 100 | 1330 |
| TFE | 20 % | 0.90 | 20.36 | 4.09 | 79.64 | 2.38 |
|  | 60 % | 1 | 00 | 1475 | 100 | 1452 |
| Ethanol | 20 % | 1.02 | 53.74 | 7.3 | 46.26 | 1.69 |
|  | 60 % | 1.4 | 24.73 | 8.8 | 75.27 | 3.82 |

### 3.5 Surface charge distribution on LUVs in presence of NSP1-CTR

The zeta potential measurement was done for the estimation of surface charge distribution of liposomes (DOPS LUV and DOPC LUV) upon addition of the NSP1-CTR peptide. The results showed that there is minor increase in the zeta potential towards the negative value for DOPS with NSP1 (−32.8 ±3.15) than DOPS alone (−28.85 ±0.21), while negligible change was found in the DOPC values **(Table 3)**. The probable reason behind this might be the charge properties of NSP1 C-terminal. The charge calculated for NSP1-CTR was −6, which thus cause electrostatic repulsion while interacting with DOPS. Ionic interactions play important role in interaction of peptide with oppositely charged lipid vesicals as reported previously (36, 53).

**Table 3.**
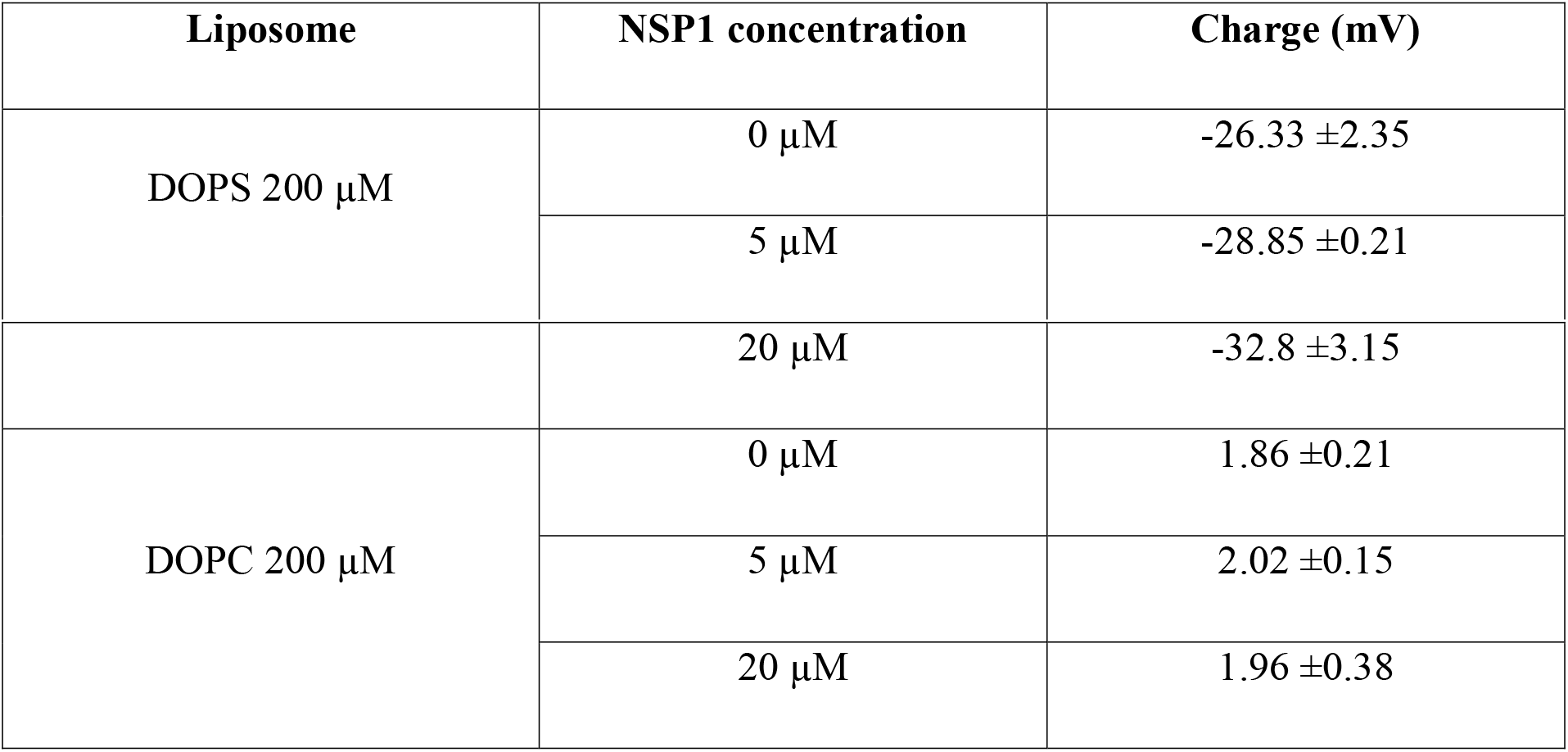
Zeta potential measurement of liposome in the presence of NSP1 C-terminal region

| Liposome | NSP1 concentration | Charge (mV) |
| --- | --- | --- |
| DOPS 200 μM | 0 μM | $-26.33 \pm 2.35$ |
| | 5 μM | $-28.85 \pm 0.21$ |
| | 20 μM | -32.8 ±3.15 |
| DOPC 200 μM | 0 μM | 1.86 ±0.21 |
| | 5 μM | 2.02 ±0.15 |
| | 20 μM | 1.96 ±0.38 |

### 3.6 Conformational dynamics of NSP1-CTR through MD simulation

We have corelated the conformational dynamics of peptide with its surrounding environment using CD spectroscopy and fluorescence spectroscopy. To complement our experiments we further performed the computational simulation. The NSP1-F1 simulation run was evidently represented the change in conformation of NSP1-CTR in presence of different environment i.e., TFE, SDS, and DOPS **(Figure 7A,B).** These changes may have occurred due to hydrophobic and electrostatic interactions that are likely to be formed during the simulation period. The gain in helix in TFE is visible with the residues 145-148 although, in presence of SDS, no such helix has appeared but there is more percentage of helix can be seen in **Figure 3.**

**Figure 7A.**
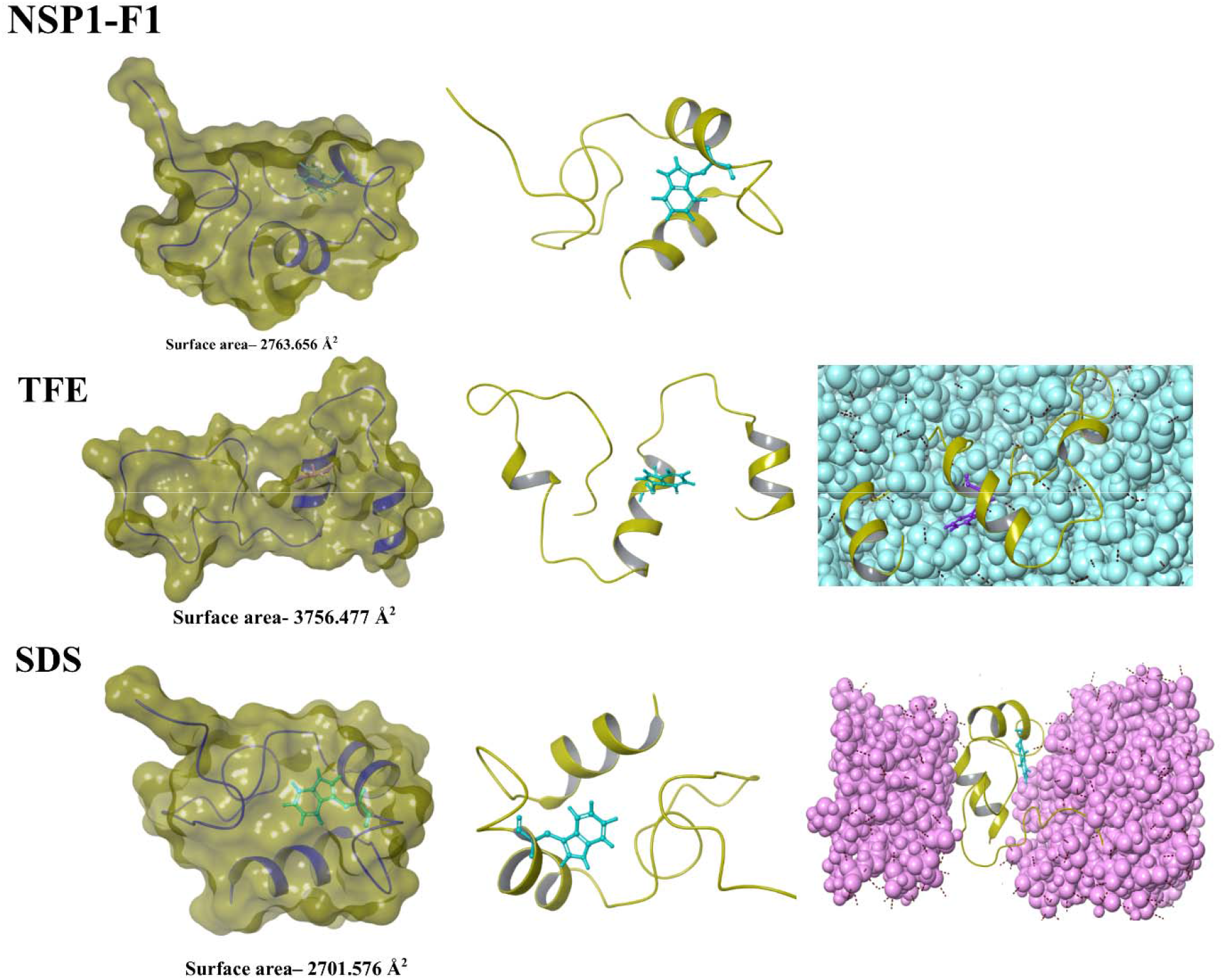
Representation of structural conformations in MD simulation highlights the change in residues near to tryptophan and surface area (measured in Schrodinger’s maestro). **Left panel:** Surface representation of final frame from simulation. **Middle panel**: Final frames from simulation. **Right panel**: TFE and SDS molecules are shown which are surrounding the structure in MD.

**Figure 7B.**
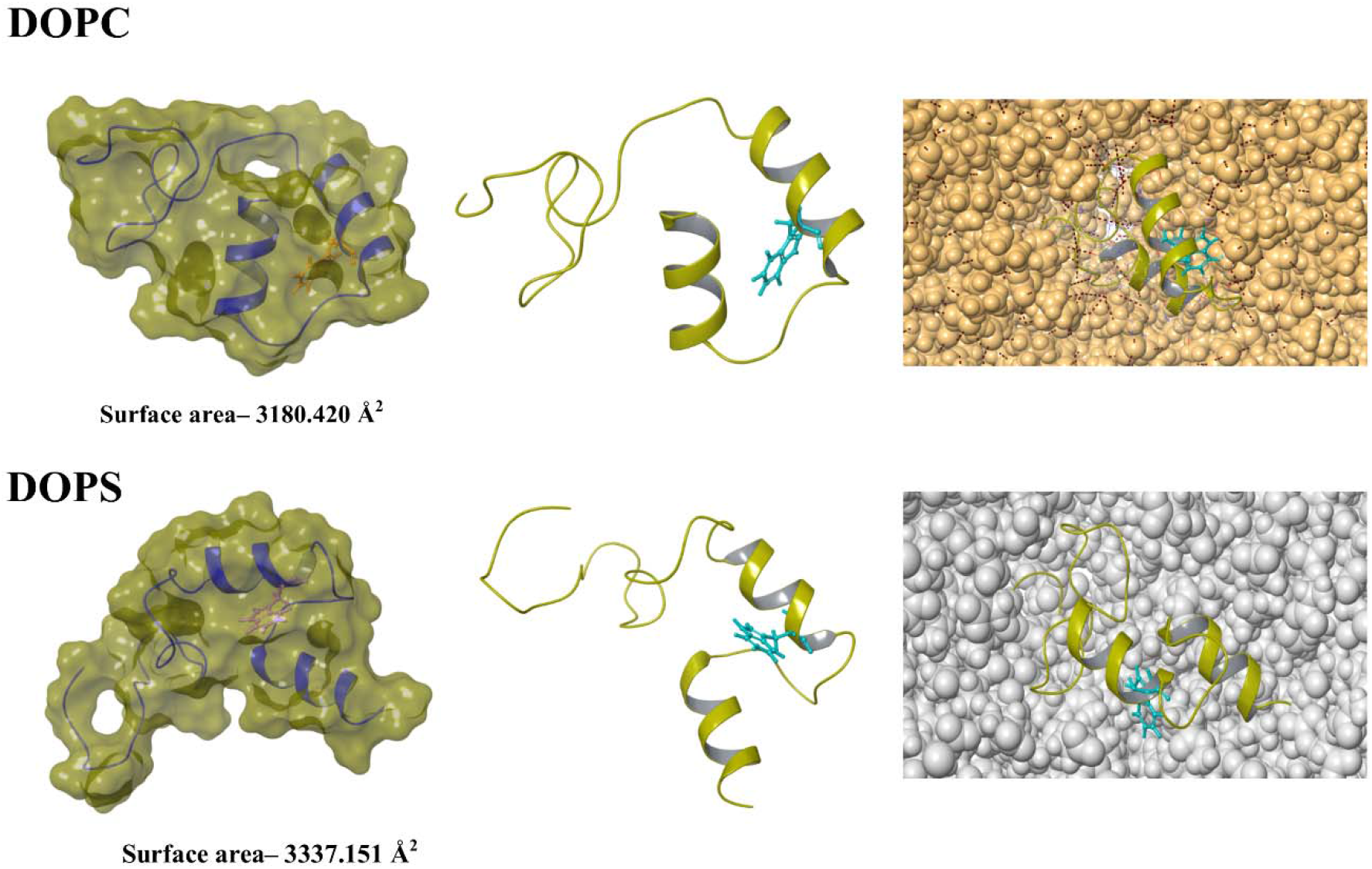
Representation of structural conformations in MD simulation highlights the change in residues near to tryptophan and surface area (measured in Schrodinger’s maestro). **Left panel:** Surface representation of final frame from simulation. **Middle panel**: Final frames from simulation. **Right panel**: DOPC and DOPS molecules are shown which are surrounding the structure in MD.

To unreveal the role of fluorophore (Trp^161^) based change in conformation dynamics, we studied the surface properties and Trp^161^ orientation in respective environment with computational simulation **(Figure 7A,B).** The surface area on NSP1-F1 in water and biological mimic environment conditions showed changes, which relate to its different conformations. The Trp^161^ has different orientation in each condition, which again represent different conformation of NSP1-CTR **(Figure 7A,B)**. The NSP1-F1 structural dynamics corroborates with our experiments and strengthen the hypothesis of interplay of multiple interactions for conformational dynamics of NSP1-CTR.

The time-dependent simulation frame analyses in **Figure 8** demonstrate these changes with RMSD, RMSF, Rg, and SASA values. First, the modeled structure of NSP1-CTR was simulated in water to check its conformation without any solvent condition, and its RMSD values found to be deviating initially (up to 0.8nm till 125ns approx.) and then stabilized at 0.6nm for next 250ns. The RMSD values remained same till 450ns, but fluctuates in last 50ns and get unstructured at some regions. These trends were also reflected in other two time-dependent parameters (Rg and SASA). According to RMSF trend, the residues at terminals fluctuates heavily but at nearby to C-terminal residues found to be less fluctuating. In the presence of TFE, the radius of gyration and RMSD values show variations in the middle of 300ns simulation. During this period, the structural transitions occurred and the region 145-148 got converted into helical conformation with one residue extra in middle helix **(Figure 8B)**. After structure gain, the RMSF values got decreased and illustrates that the structural transitions from unstructured to structured conformations are prominent. Further, in the presence of SDS, no such high fluctuations have been observed, however secondary structure analysis predicted gain of helicity **(Figure 3)**. Lastly, the CHARMM-GUI based lipid system also showed slight changes in structure as the RMSD and SASA values increased up to 100ns and then got stabilized for the rest of simulation period **(Figure 8).**

**Figure 8:**
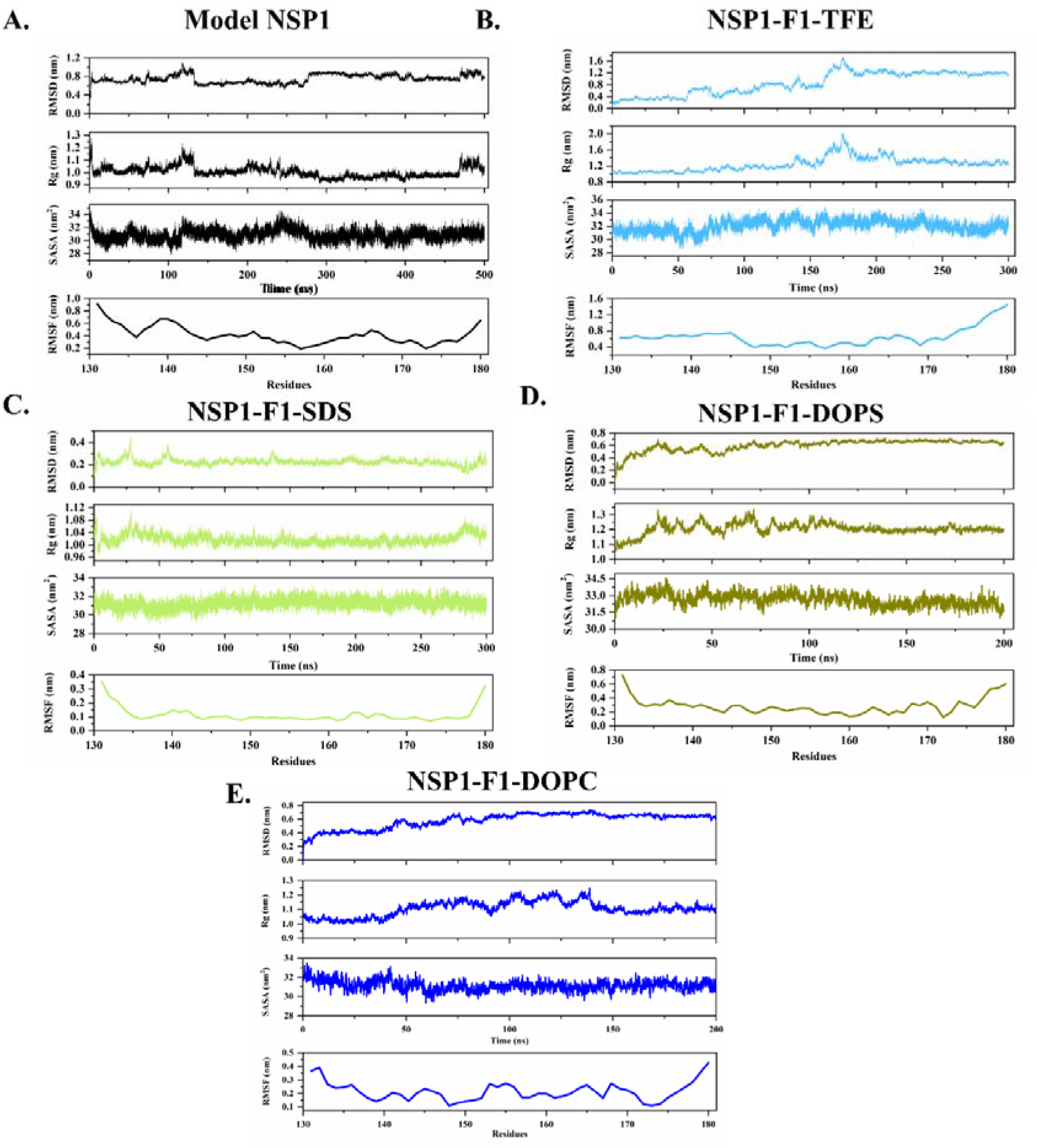
Molcular Dynamics Simulation analysis of **A.** Model NSP1, **B.** NSP1-F1-TFE, **C.** NSP1-F1-SDS, **D.** NSP1-F1-DOPS, and **E.** NSP1-F1-DOPC. From up to down RMSD, Rg, SASA, and RMSF plots are shown.

## Conclusion

In this study we have found that NSP1-CTR is an IDP in isolation. This is in-line with previous discovery using Cryo-EM that in full length (NSP1) system the C-terminal is highly flexible. Further, we investigated the folding propensity in presence of different environments and lipids and found the conformational heterogenety. The interplay of interaction forces are the driving forces, which are responsible for the protein-protein interactions and their associated functions. The binding promiscuity of IDPs provide flexibility to structural domain as well as ability to function independently with number of partner ligand. The present study allowed us to understand the NSP1-CTR, in particular how the conformation dynamics play a central role in protein interaction. The impact of hydrophobic interactions, electrostatic interaction are in agreement with the previous reports, where NSP1-CTR gain structural helicity after interaction with 40S and 80S ribosome (6). The conformational dynamics of NSP1-CTR in solution could pave a path to understand the structural ensemble of this disorder system. Further, some questions could be answered by the binding kinetics and folding studies of NSP1-CTR with ligand partner, which eventually helps us to understand the promiscuity of this peptide and design a small molecular inhibitor.

## Credit authorship contribution statement

RG and NG: study supervision. Amit Kumar and Ankur Kumar designed the experiment. Amit Kumar and Ankur Kumar conducted the experiment. Amit Kumar and PK acquisition and interpretation of computational data. Amit Kumar, Ankur Kumar, PK, and RG contributed to paper writing.

## Declaration of competing interest

All authors affirm that there are no conflicts of interest.

## Acknowledgments

All the authors would like to thank IIT Mandi for providing experimental and HPC facilities. The work was partially supported from DBT, Government of India (BT/11/IYBA/2018/06) to RG. RG is also thankful for the support from IIT Mandi. AmK was supported by DBT, Government of India (BT/11/IYBA/2018/06). PK and AnK were supported by fellowship from MHRD.

## Notes

### Competing Interest Statement

The authors have declared no competing interest.

